# A Machine Learning Framework to Predict Subcellular Morphology of Endothelial Cells for Digital Twin Generation

**DOI:** 10.1101/2022.03.21.485159

**Authors:** Miguel Contreras, Rex W. Hafenstine, William Bachman, David S. Long

## Abstract

Gaining insight into different cell behaviors is key to better understanding different pathologies. These behaviors may be explained in part through close observation of 3D cell morphology. Therefore, the objective of this research was to develop a machine learning (ML) framework that can predict 3D subcellular morphological variation of endothelial cells (ECs) to generate digital twins. ECs were cultured and their membrane, nucleus, and focal adhesion (FA) sites were stained and imaged with confocal microscopy. The multicellular confocal *z* stacks were segmented resulting in a total of 60 single-cell stacks. Fifty randomly picked cells were augmented 20-fold to train the ML framework, and the remaining 10 were used for an independent test of prediction accuracy. The ML framework was based on an open-source conditional generative adversarial network (cGAN), which was expanded to make 3D predictions using membrane only as input to predict nucleus and FA morphology. After training the framework, the results on the independent test showed an average prediction accuracy of ∼87% for nucleus and ∼70% for FA sites. The predictions were used to build a digital twin of each EC and compared to their respective ground truth, showing an average ∼79% global accuracy and ∼84% accuracy in FA-Nucleus distribution. The results presented show the effectiveness of the developed ML framework to generate digital twins of Ecs using limited amount of data. These digital twins can be used to couple EC morphology with different behaviors. The ML framework can be potentially expanded to predict morphology of other subcellular structures as well as to study other types of cells.

## 1 Introduction

Blood vessels and lymphatic vessels are lined with a monolayer of endothelial cells (ECs) called the endothelium. Endothelial cell morphology changes depending on where the cell is in the vascular system or lymphatic system and the fluid sheer stress the cell is exposed to (Davies, 2000). An elongated EC morphology is found in straight sections of arteries, while a cobblestone morphology is near pro-atherogenic regions (Liang et al., 2020). Therefore, tracking EC morphology (cell geometry and size) may indicate either a healthy or diseased cell phenotype (Alizadeh et al., 2016; Lyons et al., 2016; Yin et al., 2013). These cells can influence other cells in their neighborhood as they come into contact through junctional structures. For example, individual EC morphology has been observed to change from a spindly to a cuboidal geometry in the presence of neighboring cells (Liu & Chen, 2007). Such changes in multicellular morphology can help to understand how an individual healthy or diseased cell could influence other neighboring cells. Developing methods to measure how ECs adapt to their mechanical environment has the potential to help explain their behaviors and phenotypic change, which may be explained in part through close observation of their 3D morphology (McCarron et al., 2017). As a single cell or group of cells experience stimuli, dynamic adjustments to their morphology must occur. These morphological changes are necessary to maintain proper cell adhesion to the extracellular matrix, force distributions through the cell, and cell-cell interaction.

Gaining insight into different cell behaviors is key to better understanding different pathologies. To understand how cells respond to their surroundings will require better understanding of mechanotransmission, mechanosensing, and mechanotransduction. Mechanotransmission is the process of forces being transmitted intra- and inter-cellularly. Mechanosensing is the process by which a cellular structure or a protein senses physical cues to initiate mechanotransduction. Mechanotransduction is the process of converting physical cues into a biochemical response. For example, mechanosensing can provide new knowledge on development of cardiovascular disease (Gaetani & Messina, 2020) and cancer metastasis (Reymond et al., 2013). Observing how cell morphology, cell membrane and cell nucleus, remodel to changes in the cell’s mechanical environment is critical to understanding when a cell transitions phenotype from healthy to a cardiovascular disease state (Gaetani & Messina, 2020). For cancer cells to metastasize, they must create separations between endothelial tight junctions, thus altering the external and internal structural components of the cell (Reymond et al., 2013).

Focal adhesion (FA) sites are supramolecular contact points between the cell and the extracellular matrix (ECM) (Gaetani & Messina, 2020). Understanding the shape, distribution, and size of FA sites, can help to develop better methods to understand changes in cell morphology. To maintain proper adherence between the cell and the ECM, Fas dynamically alter their shape and mechanical structure. Focal adhesions directly influence the morphology of the cell through the process of mechanosensing. The changes in mechanical forces experienced by the Fas are transmitted throughout the cell cytoskeleton via mechanotransmission, creating a feedback loop that alters the morphology of cell membrane, nucleus, and other subcellular components (Martino et al., 2018). As the morphology of the cell is adapting to maintain structural integrity, internal subcellular structures such as the nucleus are altering the process of mechanotransduction. Processes that may be altered by changes in nucleus morphology are gene expression, DNA synthesis, and calcium signaling (Prasad & Alizadeh, 2019; Ramdas et al., 2015; Roca-Cusachs et al., 2008; Tong et al., 2017; Uhler & Shivashankar, 2016; Wang et al., 2017).

Current computational mechanical modeling of Ecs typically consider groups of cells or individual cells. One model idealized EC monolayer as a multi-cell, hexagonal, 3D, viscoelastic, multi-component model with each cell having the same idealized geometry. This model was developed to explain how fluid flow direction influences the forces experienced by structural components of Ecs (Dabagh et al., 2014, 2017). Another model used 3D single cell specific modeling on bovine aortic endothelial cells to predict stress distributions in focally adhered Ecs experiencing apically applied fluid-flow (Ferko et al., 2007). While useful, these approaches neglect emergent properties such as the heterogeneous response of cells, morphological variation, cell-cell interactions, and cellular dynamics. Another aspect that is poorly understood is the interplay between 3D focal adhesion sites and 3D nucleus morphology.

For a more comprehensive understanding of the endothelium, it is necessary to develop methods to observe the changing morphology of subcellular structures and how they are affected by the change in EC morphology over time. Live-cell imaging is however limited on the number of morphological features that a researcher can image in a single cell simultaneously, since the number of simultaneous fluorescent tags is restricted by the spectrum saturation and cell health (Ounkomol et al., 2018). This limitation prevents the live study of multiple subcellular morphologies along with biochemical events. Therefore, a new method is necessary in order to study the relationship between cell morphology and cell behavior.

One method that can be used to help overcome limitations of live-cell imaging is machine learning (ML) algorithms. When developing a ML framework, it is important to realize that morphology of sub-cellular structures is dependent on the interactions between them. For example, the cytoskeleton cannot traverse entirely through the inner cell, as the nucleus prevents this from occurring. Furthermore, the position of the nucleus is not static as it depends on internal and external forces (Tkachenko et al., 2013). Therefore, the relationship between subcellular structures are complicated and non-linear (Yuan et al., 2019). Thus, it is necessary to examine specific sub-cellular structures and how they spatially and temporally interact.

Multiple ML approaches such as convolutional neural networks (CNN) (Christiansen et al., 2018; Ounkomol et al., 2018), and conditional generative adversarial networks (cGANs) (Yuan et al., 2019) have been used to predict the morphology of cells. Each of these methods have the potential to contribute to the study of cell morphology but are either limited to 2D morphology, or do not study certain subcellular structures relevant to cell behavior such as FA sites. To improve upon previous research, this work is focused on 3D nucleus and FA sites morphology prediction using only a labeled cell membrane.

Therefore, the cGAN method (Yuan et al., 2019), which was used to predict 2D subcellular structures on single cells with the nucleus and membrane as the labeled input images, will be further developed for this approach. To capture EC morphological variation, a digital twin framework will be developed. In this work, a digital twin is a digital representation of an EC that has a cell membrane and two predicted subcellular components, the nucleus and FA sites. The overall objective of this research is to develop a ML framework that can predict 3D subcellular morphology which can be potentially combined with live-cell imaging to study the relationship between EC morphology and behavior.

## 2 Methods

### 2.1 Cell culture and culture media

Human Microvascular Endothelial cells (HMEC-1, #CRL-3243, ATCC, Manassas, VA) were maintained in complete growth media. Complete growth media consisted of MCDB 131 media (#10372019, Gibco, Grand Island, NY) supplemented with 2 mM L-glutamine (#25030081, Thermo Fisher Scientific), 10% (v/v) FBS (#15000044, Thermo Fisher Scientific) and penicillin/streptomycin (100 U/ml and 100 μg/ml concentration, respectively). Cells were seeded at a density of 10,000 cells/cm^2^ onto fibronectin-coated glass coverslips (#CS-25R17, thickness 1.5, Warner Instruments, Hamden CT) (fibronectin, 20 μg/ml, #33016-015, Thermo Fisher Scientific) mounted in 6-well plates. The cells were grown to confluence in a humidified environment at 37°C with 5% CO_2_.

### 2.2 Immunofluorescence labeling

Once the cells reached confluence, the plasma membrane of live cells were stained with Wheat Germ Agglutinin (WGA) (CF633, Biotium, Fremont, CA). Cells were washed twice in Hank’s Balanced Salt Solution (HBSS, #14025076, Thermo Fisher Scientific), then incubated with WGA (5 μg/ml) for 30 minutes at 37°C, then washed twice in HBSS. Next, the cells were simultaneously fixed (4% (w/v) paraformaldehyde (PFA) in PBS) and permeabilized in Triton X-100 (0.5% (v/v), 5 min, #T9284, Sigma-Aldrich). Then post-fixed in 4% (w/v) PFA in PBS, followed by PBS wash (3 × 5 min). To image the nucleus, cells were stained with Hoechst 33258 (1:1000, #116M4139V, Sigma-Aldrich). A blocking solution comprised of goat serum (1:20, #G9023, Sigma-Aldrich), 0.1% Triton X-100, and 0.3M glycine in 0.1% (w/v) BSA was added to cells for 30 minutes. To image FA sites, cells were blocked with the previously mentioned blocking solution for 30 minutes at room temperature. Then cells were incubated overnight with anti-paxillin (1:500, Abcam, #ab32084). This was followed by a 2-hour incubation with secondary antibody goat anti-rabbit Alexa Fluor 488 (1:500, #ab150077, Abcam, Waltham, MA, USA), and a 0.1% (w/v) BSA wash (3 × 5 min). The fibronectin coated cover slips were then removed from the well plates and mounted cell-side-down onto individual glass microscope slides using ProLong Glass (#P36982, Thermo Fisher Scientific).

### 2.3 Microscopy and image acquisition

A Leica TCS SP5 confocal microscope with a 63×/1.40 NA oil immersion lens was used to image the monolayer. UV 405 nm (nucleus), UV 488 nm (FA sites), and a HeNe 633 nm (membrane) lasers were used to sequentially excite samples. Images were acquired at 2048 pixels × 2048 pixels, with an *x*-*y* spatial resolution of 0.087 μm/pixel and *z* spatial resolution within the range of 0.1-0.15 μm/slice for different imaging sessions. A total of 8 fields of view (*i*.*e*., 8 image stacks) were acquired from cells belonging to all well plates.

### 2.4 Image processing and data augmentation

For each field of view, an overlay of the red color channel for membrane, the blue color channel for nucleus, and the green color channel for FA sites was obtained and processed as an RGB stack. From each stack, individual cells were manually segmented using *ImageJ* (ver1.52i) (Schneider et al., 2012). A total of 60 single-cell stacks were obtained from all 8 fields of view. Each stack was resized to 256 pixels × 256 pixels and all stacks were normalized to have 64 slices where if a stack had less than 64 slices it was padded with blank images and if a stack had more than 64 slices then blank slices at both ends of the stacks would be removed. For the membrane color channel, a threshold was applied to drop all pixel values below 20. Similarly, a threshold was applied to the nucleus color channel to drop all pixel values below 50. Finally, the FA color channel was processed using a top hat filter with a 3 × 3 rectangular kernel. Then, 50 randomly chosen cells from the 60 processed individual-cell stacks were augmented 20-fold performing random rotation and translation. For each augmentation iteration, a random angle between 0° and 360° and a random vertical and horizontal shift between -30 pixels and 30 pixels were chosen to rotate and shift each individual cell. This augmentation resulted in a total of 1000 single-cell stacks.

### 2.5 Data transformation

The stacks were then converted to Hierarchical Data Format version 5 (HDF5) format. For the conversion, the stacks (*i*.*e*., 1000) were transformed into NumPy arrays. The pixel values were normalized from the 0-255 range to a 0-1 range. The color channels for membrane and structure to be predicted (*i*.*e*., nucleus or FA sites) were appended to a single array of dimensions (1000, 64, 256, 256, 3) labeled as the input *X*. The labels (*i*.*e*., structure to be predicted) were transformed into a one-hot encoded vector and appended to a single array of dimensions (1000, 10) labeled as the condition *Y*, where stacks where evenly assigned to each of the 10 classes (*i*.*e*., each class consisted of 100 stacks). The color channel for cell membrane structure was separated from the images and appended to a single array of dimensions (1000, 64, 256, 256, 1) labeled as the reference *R*. Finally, the three arrays were stored into a single HDF5 file.

### 2.6 Neural network architecture and training

An open-source neural network (Yuan et al., 2019) originally developed for 2D subcellular structure prediction was extended to 3D and trained to predict the nucleus and FA sites, using only cell membrane as input. The network was based on a conditional generative adversarial network (cGAN) architecture, consisting of a generator and discriminator and where the condition *Y* determined the type of structure to be predicted, the input *X* was used to extract the ground truth structure of interest and membrane, and the reference *R* was used to extract only the membrane that would be used to generate a prediction of the structure of interest. The network was modified to make 3D predictions by replacing all convolutional layers in both the generator and the discriminator to perform 3D convolution. The loss function used for training the discriminator was a binary cross entropy, while the loss function for the generator was a sum of the binary cross entropy and a Pearson correlation metric. Two different training sessions were performed: one to predict nucleus and another one to predict FA sites. The network was trained using the augmented dataset which was split into 900 cells for training and 100 for concurrent testing. The remaining 10 cells out of the original 60 were used for an independent test of prediction accuracy at the end of training. The parameters used to train the network for each session were a learning rate of 2 × 10^−3^ for the generator and 2 × 10^−4^ for the discriminator and a batch size of 1. Both training sessions lasted 50 epochs as the network converged at this point.

### 2.7 Evaluation of predictions

To evaluate the accuracy predictions in the test set, two different metrics were used: Pearson Correlation Coefficient (PCC) for nucleus and Discrete Protein Metric (DPM) (Contreras et al., 2021) for FA sites. For nucleus, PCC was chosen since previous studies performing nucleus morphology prediction (Christiansen et al., 2018) have used the same metric for evaluation of the results. For FA sites, DPM was chosen as this metric was specifically developed to evaluate FA predictions. Before computing the DPM, both ground truth and predicted FA sites were processed using the following steps: 1) a top-hat filter with a 3 × 3 rectangular kernel was applied to the raw color channel, 2) the channel was then binarized to calculate the area of each FA, 3) only Fas that had an area ranging from 1-4 µm^2^ were kept as this range has been seen in previous studies (Kim & Wirtz, 2013), 4) the resulting binary stack was projected onto a 2D image to apply it as a mask to the original raw color channel. The DPM calculates the difference between predicted and ground truth FA sites by measuring three characteristics: distribution (*d*), shape/size (*s*), and angle of orientation (*a*) of each FA site. Equation (1) defines the calculation of the overall DPM metric,

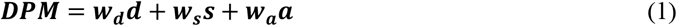

where *w_d_*, *w*_*s*_, and *w_a_* are the relative weights for FA site distribution, shape/size, and orientation angle respectively. These weights are assigned according to the relative importance of each component for the research question of interest. The sum of these weights must always equal one. For the purpose of this study, all measurements were given the same importance (*i*.*e*., all weights were set equal to 0.33). The distribution (*d*) measurement is calculated using a *k*-means clustering algorithm that divides the FA sites into five clusters and then calculates the difference in number of FA sites on each cluster in the prediction and ground truth. The shape/size (*s*) measurement is calculated by pairing each FA site in the prediction to its nearest neighbor in the ground truth and calculating the overlap using an F1 score of the bounding boxes of the predicted and ground truth FA site. The angle of orientation (*a*) measurement is calculated by once again pairing each FA site in the prediction to its nearest neighbor in the ground truth and measuring the angle with respect to the horizontal axis of the bounding boxes of the predicted and ground truth FA site to calculate their difference. The DPM was calculated on the 2D projections obtained at the end of the processing step.

A schematic of the ML training and validation workflow can be seen in Fig. 1.

**Fig. 1:**
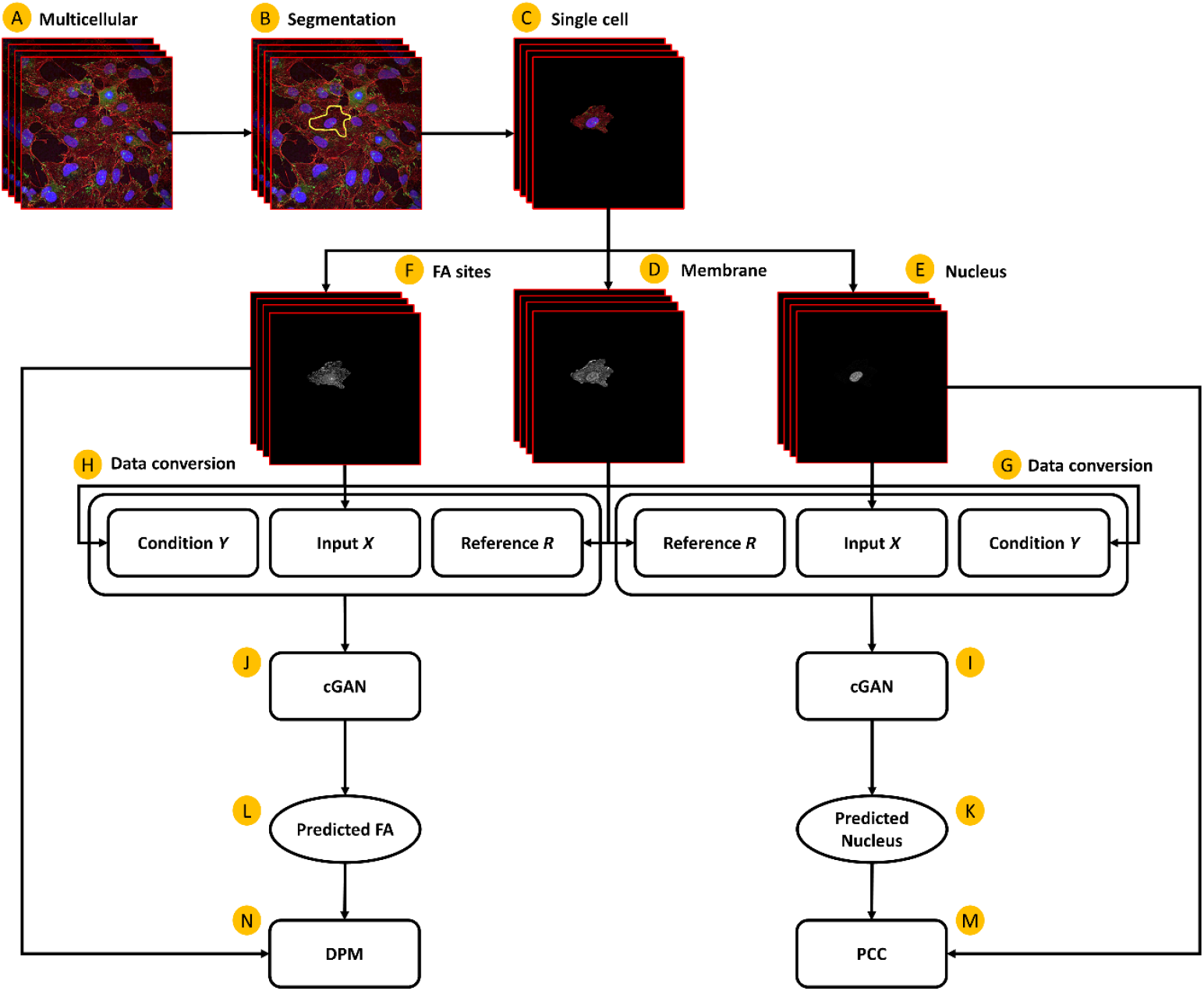
Schematic of ML training and validation: (A) Multicellular stack obtained from the confocal microscope; (B) Segmentation of single cells from cell cluster; (C) Single cell stack obtained after segmentation; (D) Extraction of cell membrane color channel into single stack; (E) Extraction of nucleus color channel into single stack; (F) Extraction of FA site color channel into single stack; (G) Conversion of image data into condition *Y* one-hot encoded vector representing the nucleus structure, input *X* array containing ground truth nucleus and membrane, and reference *R* containing membrane only; (H) Conversion of image data into condition *Y* one-hot encoded vector representing the FA sites structure, input *X* array containing ground truth FA sites and membrane, and reference *R* containing membrane only; (I) Instance of cGAN algorithm trained to predict nucleus; (J) Instance of cGAN algorithm trained to predict FA sites; (K) Predicted nucleus generated from cGAN (I); (L) Predicted FA sites generated from cGAN (J); (M) Computation of PCC metric between predicted nucleus (K) and ground truth nucleus (E); (N) Computation of the DPM metric between predicted FA sites (L) and ground truth FA sites (F).

### 2.8 Digital twin generation

Once the cGAN was fully trained for both FA and nucleus predictions, digital twins of ECs were generated using only the labeled membrane as input to the network. These digital twins were compared to the true cell (*i*.*e*., ground truth) by measuring the global accuracy score defined in Equation (2),

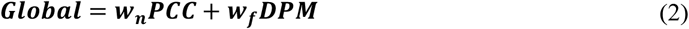

where *w*_*n*_ and *w*_*f*_ are the relative weights for nucleus and FA sites prediction importance, and PCC and DPM are the accuracy scores calculated for nucleus and FA sites predictions. The weights are assigned according to the research question of interest. For the purpose of this study, nucleus and FA sites predictions were given the same importance (*i*.*e*., the weights were set to 0.5).

For further evaluation of the digital twin similarity to its respective true cell, an FA-Nucleus distribution (*FA*-*N*_*D*_) metric, modified from a previous study (Buskermolen et al., 2018), was calculated using Equation (3),

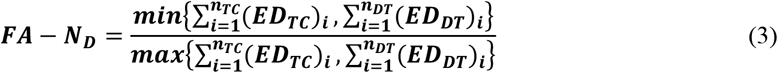

where (*ED*_*TC*_)_*i*_ and (*ED*_*DT*_)_*i*_ are the Euclidian distance between the centroids of the nucleus and the centroid of the *i*^th^ FA site in the true cell and digital twin respectively, and *n*_*TC*_ and *n*_*DT*_ are the number of FA sites in the true cell and digital twin respectively.

A schematic of the digital twin generation process can be seen in Fig. 2.

**Fig. 2:**
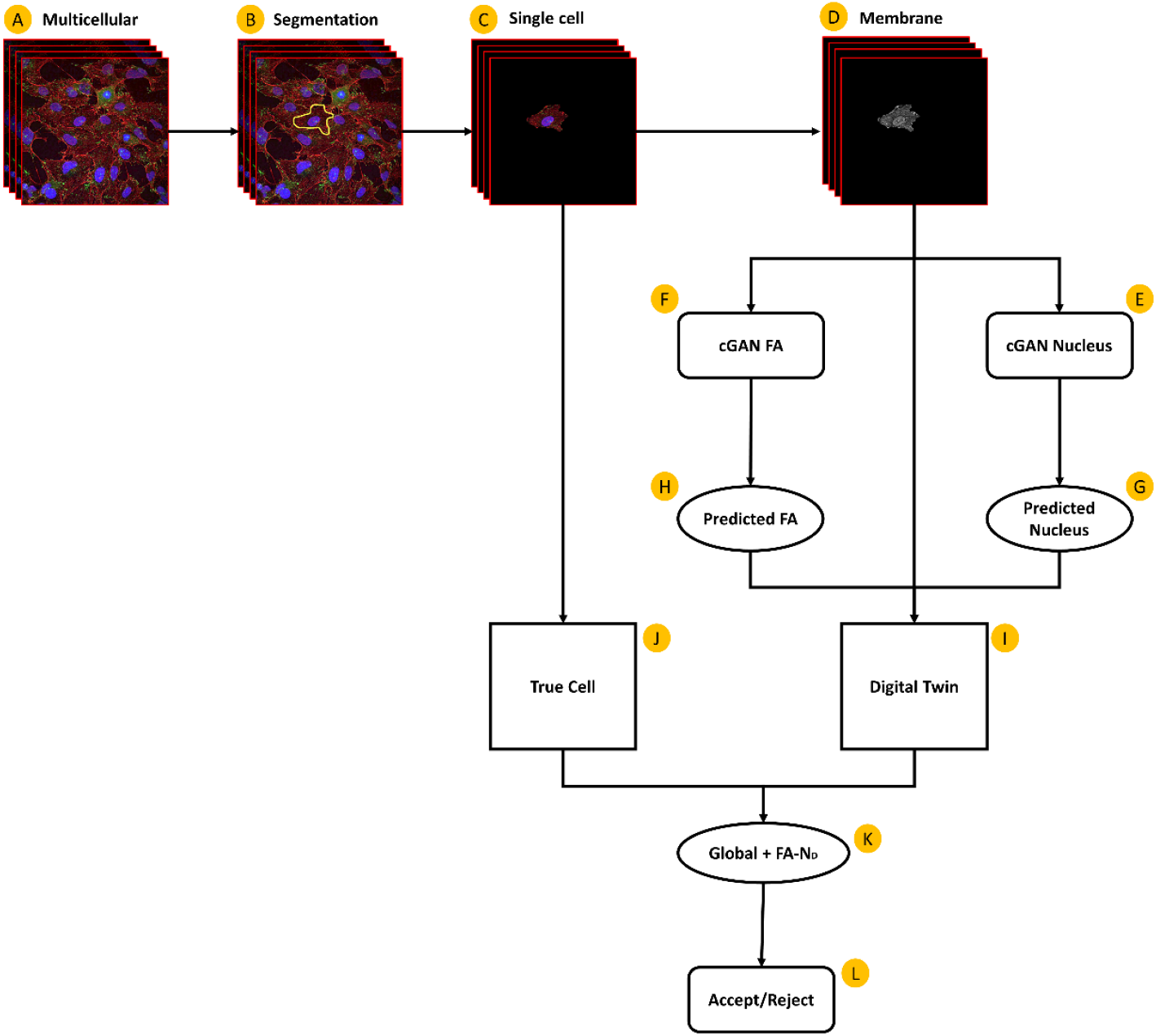
Cellular digital twin generation workflow: (A) Raw multicellular stack obtained from the confocal microscope with membrane (red), nucleus (blue), and FA sites (green); (B) Segmentation of single cells from cell cluster; (C) Single cell stack obtained after segmentation; (D) Extraction of cell membrane color channel into single stack; (E) cGAN trained for nucleus prediction; (F) cGAN trained for FA sites prediction; (G) Predicted nucleus based on membrane input; (H) Predicted FA sites based on membrane input; (I) Digital twin constructed from predicted nucleus (G), predicted FA sites (H), and cell membrane (D); (J) True cell containing ground truth nucleus and FA sites from segmented cell (C); (K) Comparison between digital twin (I) and true cell (J) using Global score and FA-N_D_; (L) Accept or reject digital twin based on criteria used to establish acceptable accuracy.

## 3 Results

The ML framework was trained using an augmented dataset of cultured and stained single ECs to predict 3D morphology of nucleus and FA sites based only on membrane morphology. The performance of the framework was then evaluated by calculating the prediction accuracy of nucleus and FA sites using an independent test containing 10 cells. Digital twins were then generated for all 10 cells and their accuracy was calculated.

The prediction accuracy for all 10 cells in the independent test set is summarized in Table 1. The average accuracy is presented as mean ± standard deviation. The average prediction accuracy for nucleus was 87.7% ± 5.1% as measured using PCC. For FA sites, the average prediction accuracy was 70.5% ± 4.6% as measured using DPM, where average *d* was 73.1% ± 10.3%, average *s* was 71.0% ± 4.6%, and average *a* was 69.6% ± 4.8%. The average global score accuracy for the independent test was 79.1% ± 3.1%. The average *FA*-*N*_*D*_ score was 84.1% ± 9.4%.

**Table 1.**
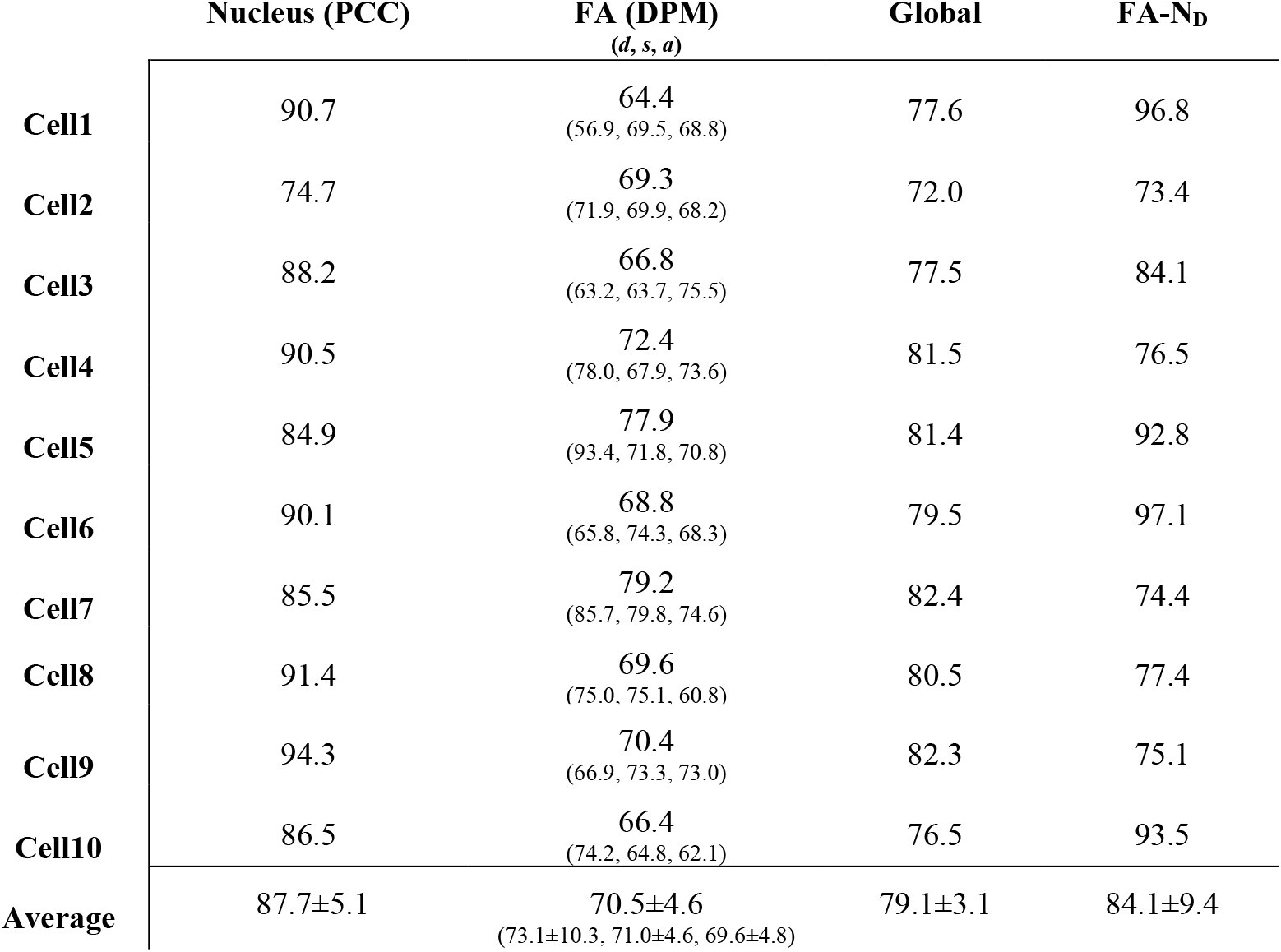
Prediction accuracy percentages for nucleus, FA sites, global score, and FA-Nucleus distribution for all cells in the independent test dataset

| | Nucleus (PCC) | FA (DPM)<br>($d, s, a$) | Global | FA-$N_D$ |
| --- | --- | --- | --- | --- |
| Cell1 | 90.7 | 64.4<br>(56.9, 69.5, 68.8) | 77.6 | 96.8 |
| Cell2 | 74.7 | 69.3<br>(71.9, 69.9, 68.2) | 72.0 | 73.4 |
| Cell3 | 88.2 | 66.8<br>(63.2, 63.7, 75.5) | 77.5 | 84.1 |
| Cell4 | 90.5 | 72.4<br>(78.0, 67.9, 73.6) | 81.5 | 76.5 |
| Cell5 | 84.9 | 77.9<br>(93.4, 71.8, 70.8) | 81.4 | 92.8 |
| Cell6 | 90.1 | 68.8<br>(65.8, 74.3, 68.3) | 79.5 | 97.1 |
| Cell7 | 85.5 | 79.2<br>(85.7, 79.8, 74.6) | 82.4 | 74.4 |
| Cell8 | 91.4 | 69.6<br>(75.0, 75.1, 60.8) | 80.5 | 77.4 |
| Cell9 | 94.3 | 70.4<br>(66.9, 73.3, 73.0) | 82.3 | 75.1 |
| Cell10 | 86.5 | 66.4<br>(74.2, 64.8, 62.1) | 76.5 | 93.5 |
| Average | $87.7 \pm 5.1$ | $70.5 \pm 4.6$<br>(73.1±10.3, 71.0±4.6, 69.6±4.8) | $79.1 \pm 3.1$ | $84.1 \pm 9.4$ |

From the 10 samples in the independent test set, Cell 5 was randomly chosen to demonstrate the power of the ML framework to generate a digital twin representative of the true cell. Fig. 3 shows a qualitative comparison of nucleus ground truth and prediction along the *z*-stack to demonstrate the accuracy (84.9% PCC) of the 3D prediction. Similarly, Fig. 4 shows a qualitative comparison of FA sites ground truth and prediction along the *z*-stack to demonstrate the accuracy (77.9% DPM) of the 3D prediction. Finally, Fig. 5 shows a visualization of the 3D structure of the digital twin compared to the true cell using AGAVE (https://github.com/allen-cell-animated/agave/releases/tag/v1.2.4). The comparison shows a qualitative validation of the global accuracy (81.4%) and subcellular distribution (92.8%) of the digital twin.

**Fig. 3:**
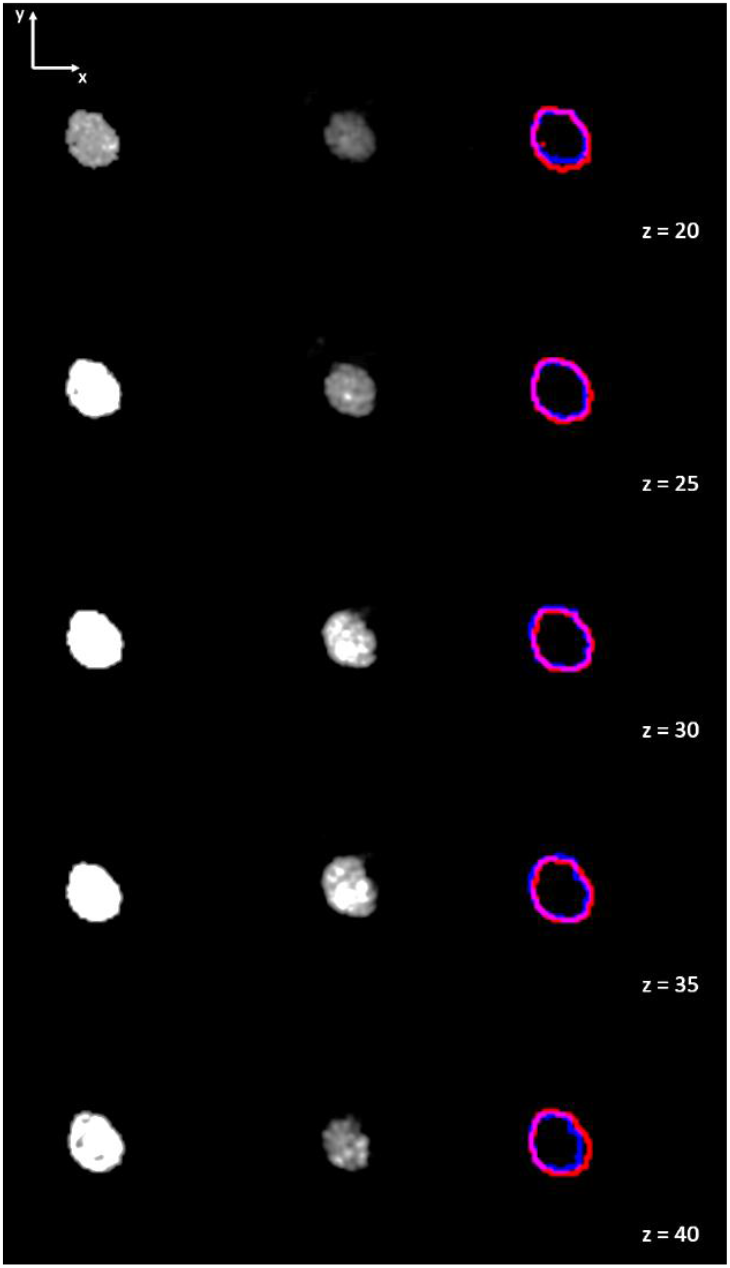
Comparison of nucleus ground truth and predictions for representative *z*-slices for Cell 5. First column represents the ground truth, second column the prediction, and third column an overlay of ground truth (red) outlines and prediction (blue) outlines. Each row represents a different slice, as indicated by the *z* slice number on the lower right corner of each row.

**Fig. 4:**
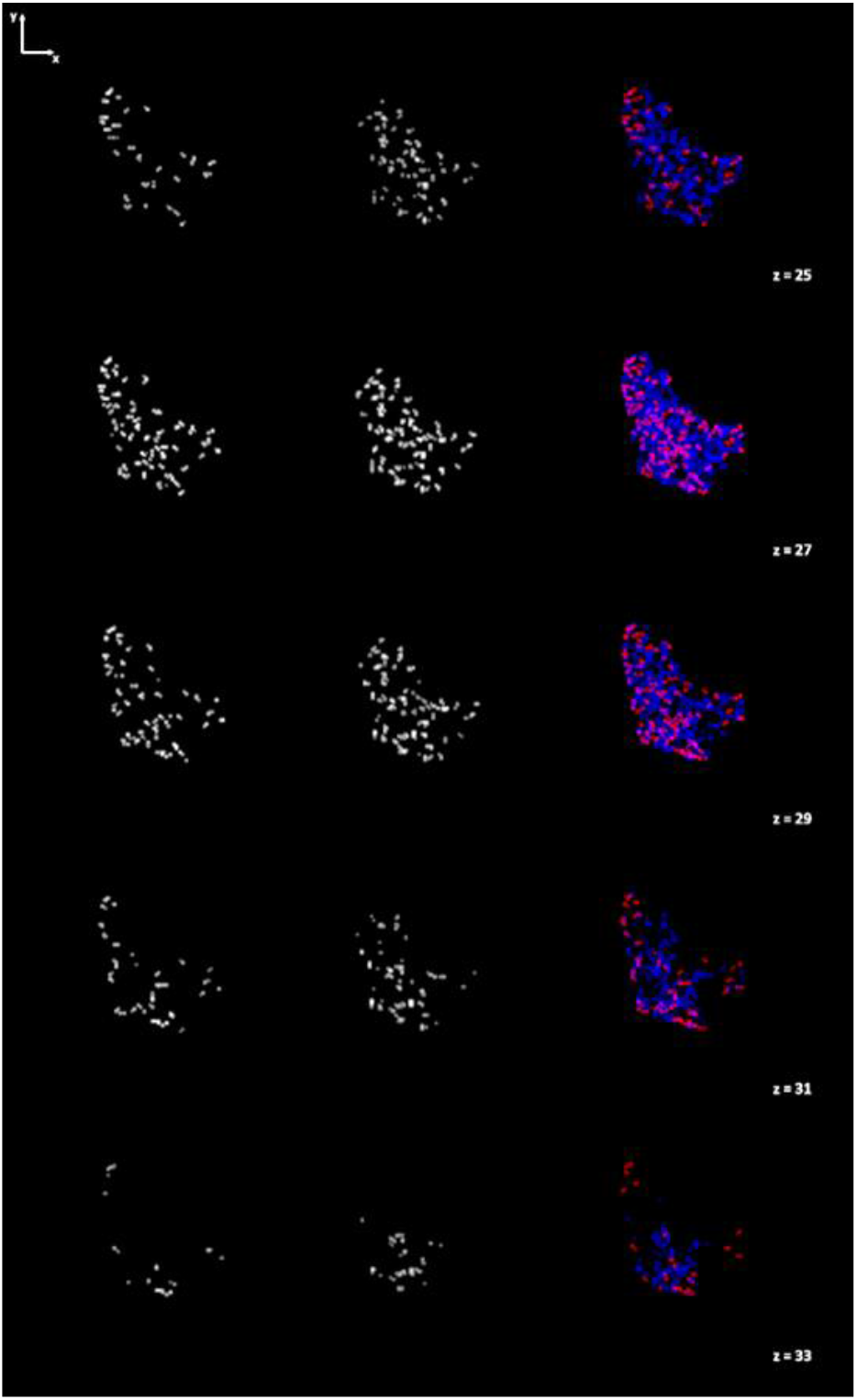
Comparison of FA sites ground truth and predictions for representative z-slices for Cell 5. First column represents the ground truth, second column the prediction, and third column an overlay of ground truth (red) and prediction (blue). Each row represents a different slice, as indicated by the *z* slice number on the lower right corner of each row.

**Fig. 5:**
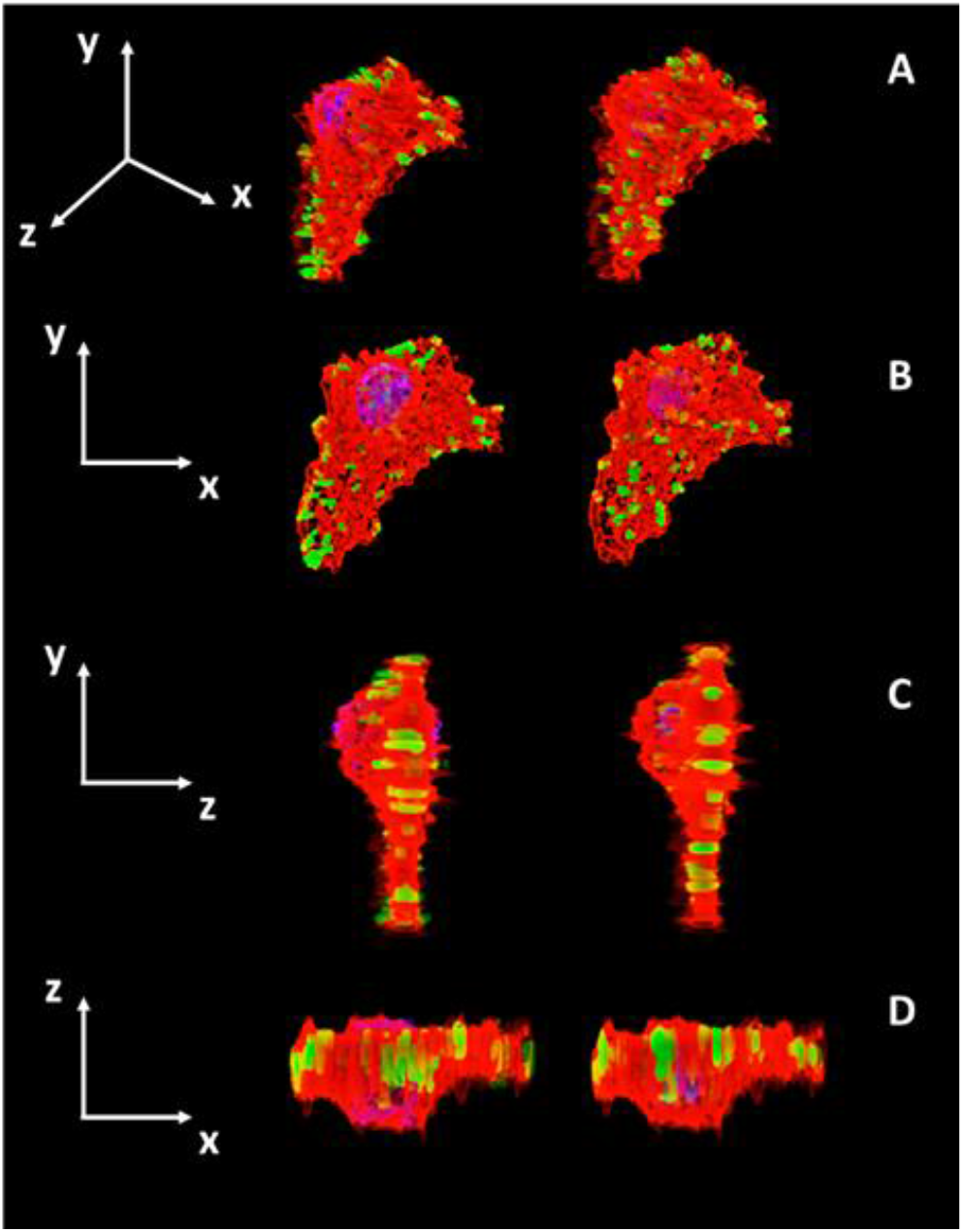
Cell 5 digital twin (right) versus true cell (left) comparison. The structures represented are membrane (red), nucleus (blue), and FA sites (green). Orthographic projection was used for visual comparison: (A) 3D representation; (B) 2D orthographic projection of front side; (C) 2D orthographic projection of right side; (D) 2D orthographic projection of top side.

## 4 Discussion

The results presented show the effectiveness of the developed ML framework to generate digital twins of ECs using limited amount of data. This framework was developed by modifying an open-source cGAN (Yuan et al., 2019) that was originally used to predict subcellular protein 2D structure from membrane and nucleus input. The developed framework can predict, with high accuracy, 3D morphology of nucleus and FA sites based only on the cell membrane morphology.

In terms of nucleus predictions, the ML framework achieves accuracy (∼87%) comparable to other methods that have predicted 2D nucleus (Christiansen et al., 2018) and higher than existing methods predicting 3D nucleus (Ounkomol et al., 2018). The fact that the proposed method used single-cell stacks might be one reason the ML framework achieves higher accuracy compared to previous methods which used multicellular stacks (Ounkomol et al., 2018). On the other hand, predictions were based on 3D membrane morphology (*i*.*e*., cell membrane fluorescence stacks) rather than on 3D whole-cell morphology (*i*.*e*., transmitted light stacks) (Ounkomol et al., 2018) which could indicate that membrane is a better predictor of nucleus morphology than the whole cell. Membrane morphology was observed to have a ‘bump’ where the nucleus was located as seen in Fig. 5C. This morphological feature of the membrane might help predict nucleus morphology and could therefore be a key reason for the increased nucleus prediction accuracy.

In terms of FA sites predictions, there are no current benchmarks for prediction accuracy of this structure. To the best of our knowledge, this study is the first one to predict FA sites. To determine if prediction accuracy is high enough, different FA morphological features were studied using the DPM metric (Contreras et al., 2021). The distribution (*d*) of FA sites is closely related to intracellular stress (Mullen et al., 2014). Therefore, a significant difference in distribution of FA sites can result in a very different mechanical behavior of a cell. The average *d* prediction accuracy achieved (∼73%) can be considered high, particularly when looking at the qualitative assessment provided in Fig. 4, where the overall distribution of FA sites in both ground truth and prediction looks similar to each other in terms of location and number of FA sites. On the other hand, size/shape (*s*) of FA sites can also be informative of the mechanical behavior of a cell. For example, the higher the substrate stiffness, the larger the size of FAs and the more force they will likely produce (Haase et al., 2014). The average *s* prediction accuracy achieved (∼71%) can also be seen to reflect the qualitative assessment from Fig. 4, where FA sites can be seen to have approximately the same area and aspect ratio in both ground truth and prediction. Furthermore, FA site orientation angle (*a*) can be related to the direction of movement of a cell. For example, a large difference in FA site angle can mean a drastic change in the direction of forces in a cell due to a change in the direction of flow and shear stress in the cell (Davies et al., 1994). The average *a* prediction accuracy achieved (∼69%) is consistent with the qualitative assessment in Fig. 4, where FA sites orientation looks similar in ground truth and prediction. Overall, the average DPM score (∼70%) reflects the high accuracy in distribution, shape/size, and angle of orientation. More importantly, as seen in the results, the average scores for the three measurements are very similar to each other which demonstrates that the ML framework can consistently predict the different morphological features of FA sites.

The generation of digital twins reflected the overall subcellular similarity between ground truth and predicted morphology. Two methodologies were used to measure this similarity. The first method, global score, was a weighted sum of the nucleus and FA sites accuracies. The weights for each structure can be assigned depending on the research question of interest. For example, if the objective of the study is to observe the relationship between gene expression and morphology, predicting nucleus morphology correctly might be more important than correctly predicting FA morphology. For this study, the same weight was assigned to both structures since the objective was limited to comparing the overall morphology of the true cell and the digital twin. The average global score (∼79%) is consistent with the qualitative assessment in Fig. 5. Nucleus in both true cell and digital twin can be seen to be at roughly the same location and have the same area and aspect ratio. Similarly, FA sites can be seen to have similar distribution and size in both true cell and digital twin. The second method to measure subcellular similarity was a FA-Nucleus distribution (*FA*-*N*_*D*_) metric. This metric measured the sum of the Euclidean distances between the nucleus centroid and FA sites centroids in both the true cell and digital twin. This measurement reflects the difference in overall subcellular distribution, which accounts for the difference in location of nucleus and FA sites. Furthermore, using the sum of these distances allows to account for differences in the number of FA sites in true cell and digital twin. The average *FA-N*_*D*_ (∼84%) is consistent with the subcellular distribution similarity seen in Fig. 5.

The digital twins generated with the ML framework can be potentially used for further studies. For example, the relationship between 3D morphology of membrane, nucleus, and FA sites could be further explored. The framework could also be expanded to predict other subcellular structures that can be included into the digital twin. Incorporating F-actin cytoskeleton and actomyosin fibers, substructures affected by changes in cellular morphology (Maniotis et al., 1997), could provide further insight into the mechanical structure of the EC. It has also been shown that biochemical signaling, such as calcium signaling, changes as a result of morphological changes of the cell and the nucleus (Prasad & Alizadeh, 2019; Tong et al., 2017). Therefore, the ML framework can potentially couple morphology with different mechanical behaviors and biochemical events. Furthermore, even though this framework was used on ECs, it has the potential to be expanded and applied to other cell types to study their subcellular morphology.

The methods presented in this paper have used morphology derived from fixed cells. While this can give a snapshot in time of what the EC is experiencing, it lacks dynamic behaviors. These behaviors may occur on different time scales. Therefore, future work needs to be adapted to live-cell imaging. This adaptation can be enabled by using the current ML framework. A next step can include live-cell staining the structures studied in this paper (*i*.*e*., membrane, nucleus, and FA sites) to observe changes in subcellular morphology over time. The ML framework could then be used to predict changes in FA and nucleus morphology based on changes in membrane morphology. Capturing these dynamic changes in morphology can enable coupling to biochemical signaling, for a more comprehensive understanding of how cell morphology and behavior are related.

An area that could be improved is the sample size of the experiment. Of the 60 processed images, 50 of them were augmented to create 1000 single cell stacks. Even though the results of the 3D predictions are promising, having more images that are not augmented can potentially improve the accuracy of the predictions. Inputting images that have not been manipulated will provide the ML framework with more morphological variability instead of seeing images that are only shifted/rotated. Increasing the dataset size can also help determine if membrane morphology is enough input to predict nucleus and FA morphology. From the results presented in this paper, it could be argued that membrane morphology might be enough to predict nucleus morphology. However, for FA morphology more information might be needed. Adding other subcellular structures to the input could potentially increase the accuracy of these predictions. For example, adding F-actin to the input can potentially improve FA predictions, as the morphology of this structure is closely related to FA morphology (FA sites serve as anchoring points for F-actin).

Another area that could be improved is the validation methods for the digital twins. Global and *FA-N*_*D*_ scores provide great insight into the similarity between true cell and digital twin, but they each have limitations. Global score provides a measurement that accounts for both location and shape/size of nucleus and FA sites. However, the accuracy score is calculated independently for nucleus and FA sites and, therefore, does not account for the relationship between these two structures. On the other hand, *FA-N*_*D*_ provides an accuracy score based on the distribution of FA sites with respect to the nucleus, thus providing a measurement of the relationship between them. However, *FA-N*_*D*_ does not account for shape/size of nucleus and FA sites. A combination of these two scores can be used to validate digital twins based on an established acceptable accuracy threshold. To determine this threshold, other tests might be needed. For example, mechanical simulations of the whole cell can be a great resource. By running these simulations on both true cells and digital twins, differences between their mechanical behavior can be used to determine which accuracy level is acceptable to validate the digital twins.

As the ML framework is further developed, more subcellular structures can be predicted and biochemical signaling can be integrated into the digital twin to create an interactive 3D model. This would differ from the static 3D digital representation of the cell currently presented. It is the vision of the authors to create an interactive 3D digital twin that involves a cluster of ECs which will create opportunities to view dynamic spatial and temporal changes over multiple cells. Elements of the EC that could be explained with the interactive 3D model are how ECs change their phenotype to respond to mechanical stimulus, how an EC transitions from a healthy state to a diseased state, how do ECs respond to changes of the extracellular matrix stiffness, how are biochemical signals transmitted through the EC cluster, and how do multiple subcellular structures remodel to adjust to changes in the mechanical and biochemical environment.

## 5 Conclusion

The data presented demonstrates that the ML framework can predict the 3D nucleus and focal adhesion sites from only membrane stacks. These predictions were used to generate cellular digital twins. These digital twins can be applied to couple individual behaviors on live cells, such as mechanosensing, and detailed 3D subcellular morphology of the same cells to identify and predict emergent phenotypes.

## 6 Conflict of Interest

The authors declare that the research was conducted in the absence of any commercial or financial relationships that could be construed as a potential conflict of interest.

## 7 Author Contributions

Conceptualization, M.C., R.W.H, and D.S.L.; Methodology, M.C., W.B. and D.S.L.; Software, M.C.; Data Acquisition, W.B.; Data Analysis, M.C.; Investigation, M.C.; Writing – Original Draft, M.C., and R.W.H; Writing – Review & Editing, M.C., R.W.H., W.B., and D.S.L.; Visualization, M.C.; Supervision, D.S.L.; and Funding Acquisition, D.S.L.

## 8 Funding

Research project funded in part by the Kansas Idea Network of Biomedical Research Excellence (K-INBRE); Grant number: P20 GM103418.

## 9 Data Availability Statement

The datasets and code used for this study can be found in the Cellular Digital Twin Github Repository: https://github.com/mechanobiology/cellular-digital-twin. The repository contains the segmented single-cell stacks. To obtain the multi-cellular raw image stacks please contact the corresponding author.

## Notes

### Competing Interest Statement

The authors have declared no competing interest.

